# High variability impairs motor learning regardless of whether it affects task performance

**DOI:** 10.1101/111823

**Authors:** Marco Cardis, Maura Casadio, Rajiv Ranganathan

**Affiliations:** Department of Informatics, Bioengineering, Robotics and System Engineering, University of Genova, Genova, Italy; Department of Kinesiology, Michigan State University, East Lansing, MI, USA; Department of Mechanical Engineering, Michigan State University, East Lansing, MI, USA

## Abstract

Motor variability plays an important role in motor learning, although the exact mechanisms of how variability affects learning is not well understood. Recent evidence suggests that motor variability may have different effects on learning in redundant tasks, depending on whether it is present in the task space (where it affects task performance), or in the null space (where it has no effect on task performance). Here we examined the effect of directly introducing null and task space variability using a manipulandum during the learning of a motor task. Participants learned a bimanual shuffleboard task for 2 days, where their goal was to slide a virtual puck as close as possible towards a target. Critically, the distance traveled by the puck was determined by the sum of the left and right hand velocities, which meant that there was redundancy in the task. Participants were divided into five groups – based on both the dimension in which the variability was introduced and the amount of variability that was introduced during training. Results showed that although all groups were able to reduce error with practice, learning was affected more by the amount of variability introduced rather than the dimension in which variability was introduced. Specifically, groups with higher movement variability during practice showed larger errors at the end of practice compared to groups that had low variability during learning. These results suggest that although introducing variability can increase exploration of new solutions, this may come at a cost of decreased stability of the learned solution.

## Introduction

Motor variability plays a central role in motor learning. From its original conception as "noise" in information-processing theories, where the goal of learning was to reduce all variability (Fitts and Peterson, 1964), recent approaches such as reinforcement learning and dynamical systems theory have highlighted the adaptive value of motor variability in being able to escape suboptimal solutions, and explore new solutions (Davids et al., 2003, 2006; Stergiou et al., 2006; Thelen, 1995). As a result, there has been a renewed interest in understanding the role of motor variability during learning, and specifically if introducing variability during practice can facilitate learning in both normal and clinical populations.

However, in spite of the large number of studies that have looked at this issue, the evidence for introducing motor variability to facilitate motor learning is quite mixed. Several studies on variable practice, originally derived from ‘schema theory’ (Schmidt, 1975), showed beneficial effects of variable practice on learning and generalization to novel tasks (Catalano and Kleiner, 1984; Moxley, 1979; Wrisberg and Ragsdale, 1979; Wulf and Schmidt, 1997). However, in a large meta-analysis of these studies, these effects were shown to be less robust than originally assumed (Van Rossum, 1990). More recently, Wu et al. (Wu et al., 2014) found that variability prior to learning a task is associated with faster rates of adaptation in both error-based and reinforcement tasks. However, again, a recent analysis of several datasets showed that effects of variability on rates of adaptation were not entirely consistent with the original prediction (He et al., 2016). These results suggest that in spite of several theoretical predictions of how variability should affect learning, the experimental evidence remains rather inconclusive.

One potential solution to address this mixed evidence is not to treat all motor variability the same, but rather separate variability based on its effect on task performance (Ranganathan and Newell, 2013). Because of the redundancy present in most motor tasks, motor variability can be separated into a “task-space” component (i.e., variability that affects task performance) and a “null-space” component (variability that has no effect on task performance), and this distinction has been central to several recent techniques that have been developed to examine variability (Cohen and Sternad, 2009; Cusumano and Cesari, 2006; John et al., 2016; Muller and Sternad, 2004; Scholz and Schöner, 1999). In line with this, Singh et al. (Singh et al., 2016) found that the amount of null space variability was correlated to the rate of learning, but not task space variability. These results suggest that this distinction between the types of variability during learning may provide a basis for better understanding the effects of variability on learning – however, a direct causal test is necessary to test this hypothesis.

The purpose of this study was to examine the causal influence of directly increasing either task-space or null-space variability in the learning of a redundant task. Participants learned a bimanual shuffleboard task where participants had to throw a virtual puck to a target placed at a specified distance. Using a manipulandum, we perturbed hand velocities during the throw to increase variability during practice, and examined the learning across two days of practice. Specifically, we tested how learning was affected by (a) the amount of variability introduced, and (b) the dimension (i.e. task or null space) in which the variability was introduced.

## Methods

### Participants

50 healthy college-aged participants (ages 18-26, 32 female) with no history of neurologic or orthopedic impairments volunteered to participate in the experiment. Participants received course credit for participation. All participants except one were right-hand dominant. Participants provided written informed consent and procedures were approved by the Michigan State University IRB.

### Apparatus

Participants used a bimanual planar manipulandum (KINARM Endpoint Robot, BKIN technologies, Kingston, ON) for performing the task. The position data from the end-effector of each arm was sampled at 2000 Hz. The screen was positioned above a semi-silvered mirror so that the visual feedback to the participant was aligned at the level of the handles of the manipulandum (Fig 1A).

**Figure 1.**
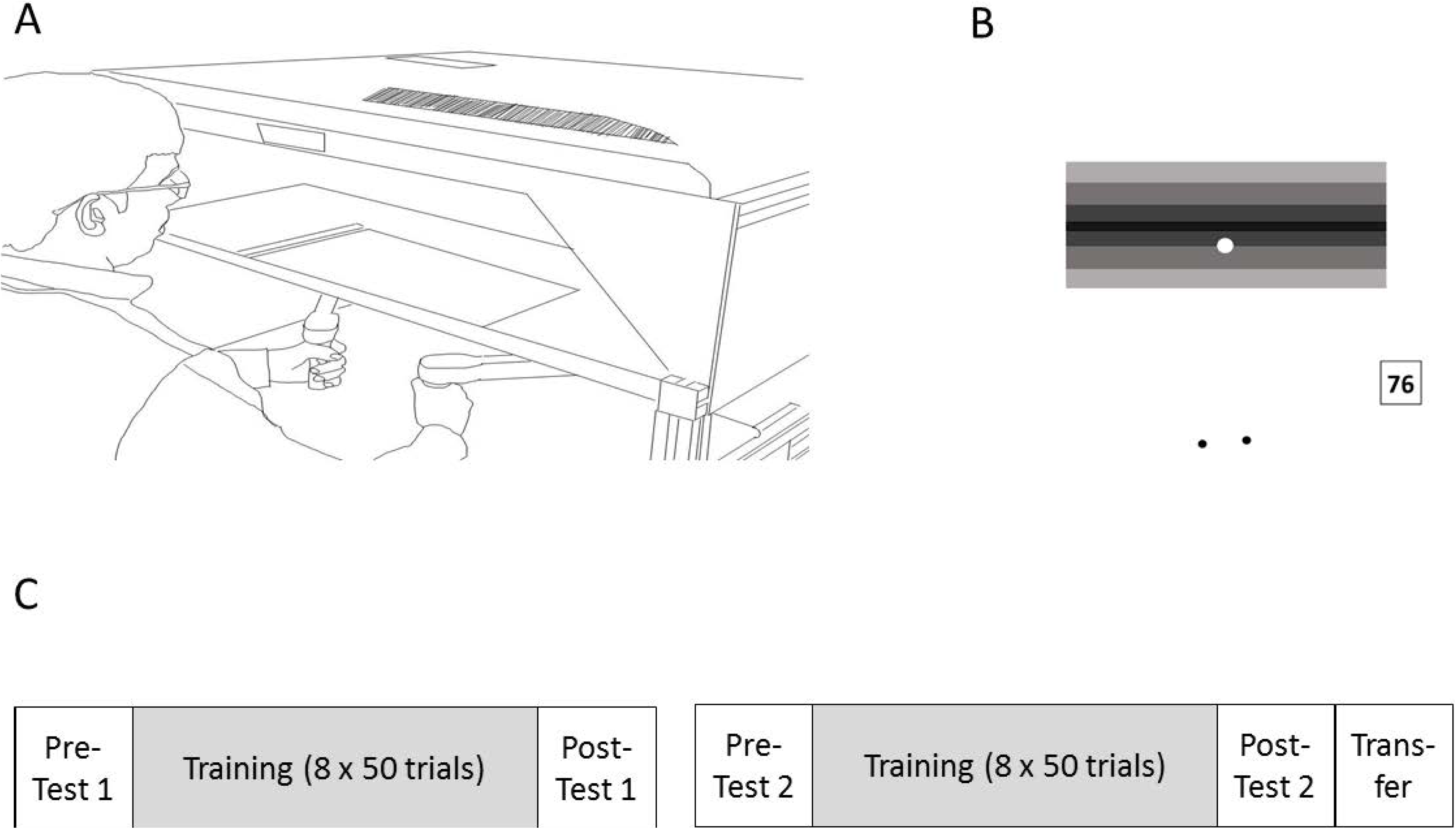
Experimental setup and protocol. (A) Participants held a bimanual planar manipulandum to perform a virtual shuffleboard task. (B) Schematic of the virtual shuffleboard task – participants attempted to land the virtual puck (white circle) close to the center target line, and received a score (from 0-100) that depended on the error. The two black dots below the target indicate the position of the hands, but were not visible to the participant. (C) Practice schedule for each day for all groups – training blocks differed between groups, whereas the test blocks (pre, post and transfer tests) were identical for all groups.

### Task

The task was a bimanual virtual shuffleboard game. Participants were instructed to hold both handles of the manipulandum and slide a virtual puck toward a target line. The goal of the participants was to release the puck so that it landed as close to the center of the target as possible (see Fig. 1B). At the beginning of each trial, participants were shown two home positions (one for each hand) and instructed to move each hand to its respective home position. Once they did, the cursor for each hand disappeared and they saw a single virtual puck on the screen whose position was computed as the average position of the two hands (Diedrichsen, 2007; Mutha and Sainburg, 2009). They were then asked to make a discrete throwing movement using both hands. Once the puck crossed a certain distance threshold from the home position (10 cm), the puck was “released” from the hand. Depending on the magnitude of the release velocity of the puck at this instant, the participant would see the puck slide a certain distance toward the target. This was a 1-D task, i.e. the direction of movement of the puck was always straight ahead toward the target (i.e. regardless of the direction of motion of the hands, the puck always traveled only straight ahead). At the end of each trial, the participant was able to see where the puck stopped relative to the target, and also received a score that depended on the absolute error (i.e. the distance between where the puck stopped and the center of the target).

Critically, the task was designed so that the magnitude of the release velocity of the puck (which determined how far the puck would slide) depended on the sum of the magnitude of the release velocities of the two hands at the instant of release (i.e. v = v_L_ + v_R_ where v is the release velocity of the puck, and v_L_ and v_R_ are the right and left hand velocities, respectively). As a result, this task was redundant - i.e. participants could use different combinations of right and left hand velocities to make the puck land in the center of the target. The solution space for this task is represented by the goal equivalent manifold (GEM) shown in Figure 2A. Specifically, the null-space represents the direction along which combinations of left and right hand velocities that yield successful task performance, and the task-space represents the direction orthogonal to the null space. This feature of the task allowed us to manipulate variability specifically along task- or null-spaces.

### Introducing null and task space variability for different groups

To introduce variability in this task, we perturbed the hand velocities during the throw by applying forces through the manipulandum. The manipulandum generated a velocity-dependent viscous field on both hands, and the strength of the viscous field on each hand was determined by constant viscosity coefficients - b_L_, b_R_ - according to the following equation:

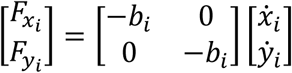

where 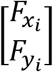 was the force field generated by the manipulandum and 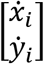 was the hand velocity vector for the *i* hand, i.e. *i*=R for the right and *i*=L for the left hand.

By altering these viscosity coefficients on each hand, we introduced variability in the task space or null space. Specifically, to introduce task space variability, we positively covaried the viscosity coefficients so that on a particular trial, both hands would either go faster, or go slower than average. In contrast, to introduce null space variability, we negatively covaried the coefficients so that on a particular trial, one hand went faster than average, whereas the other went slower than average. In addition to introducing variability along the task and null space dimensions, we also altered the amount of variability introduced by adjusting the magnitude of the changes in the viscosity coefficients (Figure 2B-E).

**Figure 2.**
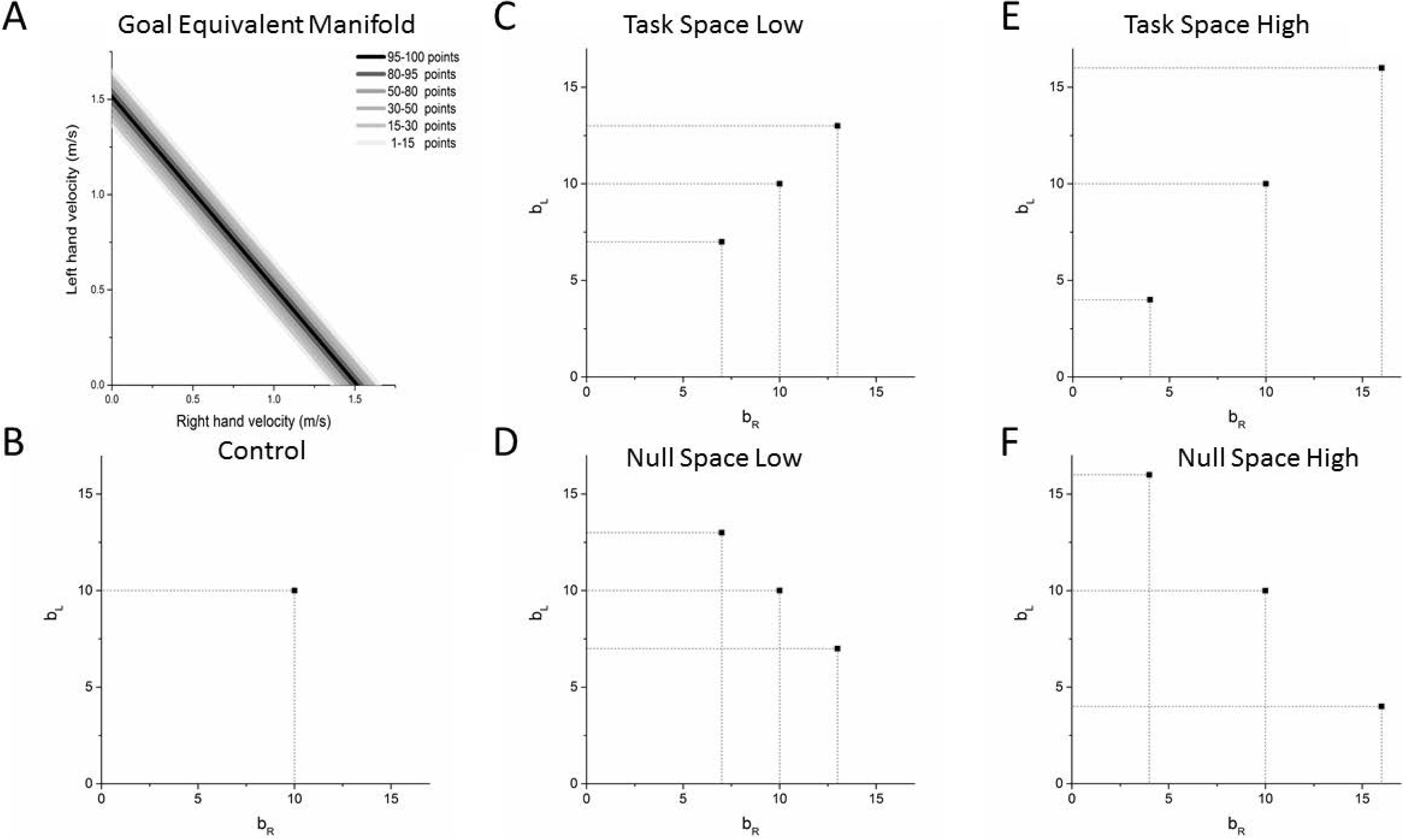
(A) Goal-equivalent manifold (GEM) for the virtual shuffleboard task. Dark colors indicate regions of better task performance (puck closer to target), whereas lighter colors indicate poorer task performance (puck farther away from target). The null space for this task is along the dark line (with −1 slope), whereas the task space is perpendicular to the null space. (B)-(E) viscosity coefficients (b_L_,b_R_) for the different groups during the training blocks designed to introduce variability of different amounts along the task and null spaces.

Based on the dimension in which the variability was introduced (task or null space) and the amount of variability introduced (high or low), participants were assigned into one of five groups as described below:

For the control group, the viscosities (b_L_, b_R_) on each trial were always set at (10,10) Ns/m and were unaltered throughout the training block. For the task space low group, to create variability along the task space, we positively covaried b_L_ and b_R_ so that the viscosities (b_L_,b_R_) on each trial were either (7,7), (10,10), or (13,13) Ns/m. For the task space high group, we positively covaried b_L_ and b_R_ similar to the task space low group, but increased the magnitude of the change so that the viscosities (b_L_,b_R_) on each trial were either (4,4), (10,10), or (16,16) Ns/m. For the null space low group, to create variability along the null space, we negatively covaried b_L_ and b_R_ so that the viscosities (b_L_,b_R_) on each trial were either (7,13), (10,10), or (13,7) Ns/m. Finally for the null space high group, we negatively covaried b_L_ and b_R_ similar to the null space low group, but increased the magnitude of the change so that the viscosities (b_L_,b_R_) on each trial were either (4,16), (10,10), or (16,4) Ns/m

In all conditions where b_L_ and b_R_ were varied, the combinations were chosen randomly from trial to trial, with the constraint that each combination had to be performed once before repeating a combination.

### Procedure

The protocol for participants across the two days is shown in Figure 1C.

#### Familiarization

In the familiarization block, participants were asked to throw the puck for 10 trials so that they could get used to performing the task with the bimanual manipulandum and understand the scoring system.

#### Pre-test

In the pre-test, the target was placed in a specific location so that participants had to release the puck at a velocity of 1.5 m/s (i.e. v_L_ + v_R_ = 1.5) to reach the center of the target. The manipulandum provided a background viscous force of 10 Ns/m for each hand during the pre-test. The pre-test consisted of 50 trials. The viscosities (b_L_,b_R_) were constant at (10, 10) Ns/m throughout all 50 trials.

#### Training blocks

In the training blocks, the participant had to release the puck at a velocity of 1.5 m/s to reach the center of the target similar to the pre-test. However, depending on the group that the participant was assigned to, the manipulandum provided trial-to-trial variations in the viscosity level to increase the variability along the task-or null-space (as described above).

#### Post-test

The post-test was identical to the pre-test. Because the pre- and post-test were identical for all groups, we used the difference between the pre-test and post-test as our metric of learning

#### Transfer test

Finally, to examine how well learning in this task was generalizable, we also performed a transfer test at the end of day 2 where the target was shifted further away so that participant had to release the puck at a higher velocity (1.72 m/s) to reach the center of the target. All other conditions were identical to the pre- and post-tests.

### Data Analysis

Task performance: The main outcome variable was the absolute error i.e. the distance between the puck and the center of the target.

Task and null space variability: Since the distance of the puck depended only on the sum of the left and right hand velocities at release, each throw can be represented as a coordinate point (vl, vr). As mentioned earlier, the solution manifold for this task is given by v_L_+ v_R_ = 1.5 m/s (for the pre-test, training, and post-test), and v_L_+v_R_ = 1.72 m/s (for the transfer test). We projected each point on to the task space and null space, and computed the variability V_task_ and V_null_ along each dimension respectively.

Aspect ratio: To examine the movement strategy, we computed the aspect ratio as the ratio of the null space to task space variability (V_null_/V_task_) (Latash et al., 2002; Scholz and Schöner, 1999). This metric allowed us to examine changes in the null space variability relative to the task space variability, and how that differed between groups, and with practice.

Lag −1 autocorrelation in task and null space: Finally, to examine the temporal structure of the variability, we computed the lag-1 autocorrelation (referred to as ACF-1) along the task- and null-spaces. Similar to the computation of variability, each point was projected on to the null and task space, and the lag-1 autocorrelation was computed on these projected points in each space separately. The lag-1 autocorrelation is a measure of how much participants learn from a movement, and correct on the subsequent movement, and is related to the learning rate (van Beers et al., 2013a; Dingwell et al., 2013)

### Statistical Analysis

At the beginning of each practice block, we observed warm-up decrement (Adams, 1952) - i.e. a temporary decline in performance for the first few trials after a rest period. Because our variability measures were sensitive to the presence of outliers, we excluded the first 10 trials in each block to minimize the warm up effects, and used only the remaining 40 trials for the analysis.

First, to ensure that the perturbations in the viscosity coefficients had the desired effect on null space and task space variability during training (i.e. the manipulation check), we compared the null and task space variabilities across groups during the first block of training using a one-way ANOVA, followed by the Dunnett’s post-hoc test that compared all other groups to the control group.

To measure differences between the groups during learning, all dependent variables were analyzed using a 5 × 5 (test × group) mixed model ANOVA. The test (pre-test1, post-test1, pretest2, post-test2, transfer) was the within-subjects factor and the group (control, task-low, task-high, null-low, null-high) was the between-subjects factor. The Greenhouse-Geisser correction was used for violations of sphericity. In post-hoc comparisons, rather than run all pairwise comparisons, we examined three separate effects: (i) all groups were compared against the control group using the Dunnett’s test. In addition, to test our hypothesis about the amount and dimension that variability was introduced, we performed two planned contrasts: (ii) the low variability groups (null space low and task space low) were compared to the high variability groups (null space high and task space high), and (iii) the null space groups (null space low and null space high) were compared to the task space groups (task space low and task space high). The significance level for all tests was set at 0.05.

## Results

When examining the results of the pre-test1, 3 out of the 50 participants had very high errors relative to the rest of the population (outliers were identified if the scores were > Q3 + 1.5*IQR, where Q3 refers to the 75^th^ percentile of data, and IQR refers to the interquartile range). These individuals were removed from further analysis: as a result the final sample size used for analysis in each of the group was as follows: control (n = 10), task space low (n = 10), task space high (n = 9), null space low (n = 9), and null space high (n = 9).

### Effects of perturbations on null and task space variability

In order to verify if the perturbations created the desired effect, we examined the task and null space variability during the first block of practice.

For the task-space variability, there was a significant main effect of group, F(4,42) = 13.17, p <. 001. Post-hoc comparisons indicated that task space low, task space high and null space high groups all had higher task space variability compared to the control group.

For the null space variability, there was also a significant main effect of group, F(4,42) = 28.663, p <.001. Post-hoc comparisons indicated that the null space low and null space high groups had significantly higher null space variability than the control group.

### Absolute Error

All participants learned how to reduce their error with practice, but the improvements differed between groups (Fig. 3A-B). There was a significant main effect of test, F(3,126) = 71.02, p <.001, that was mediated by a significant test × group interaction, F(12, 126) = 1.85, p =.047. The main effect of group was not significant, F(4,42) = 1.574, p =.199.

**Figure 3.**
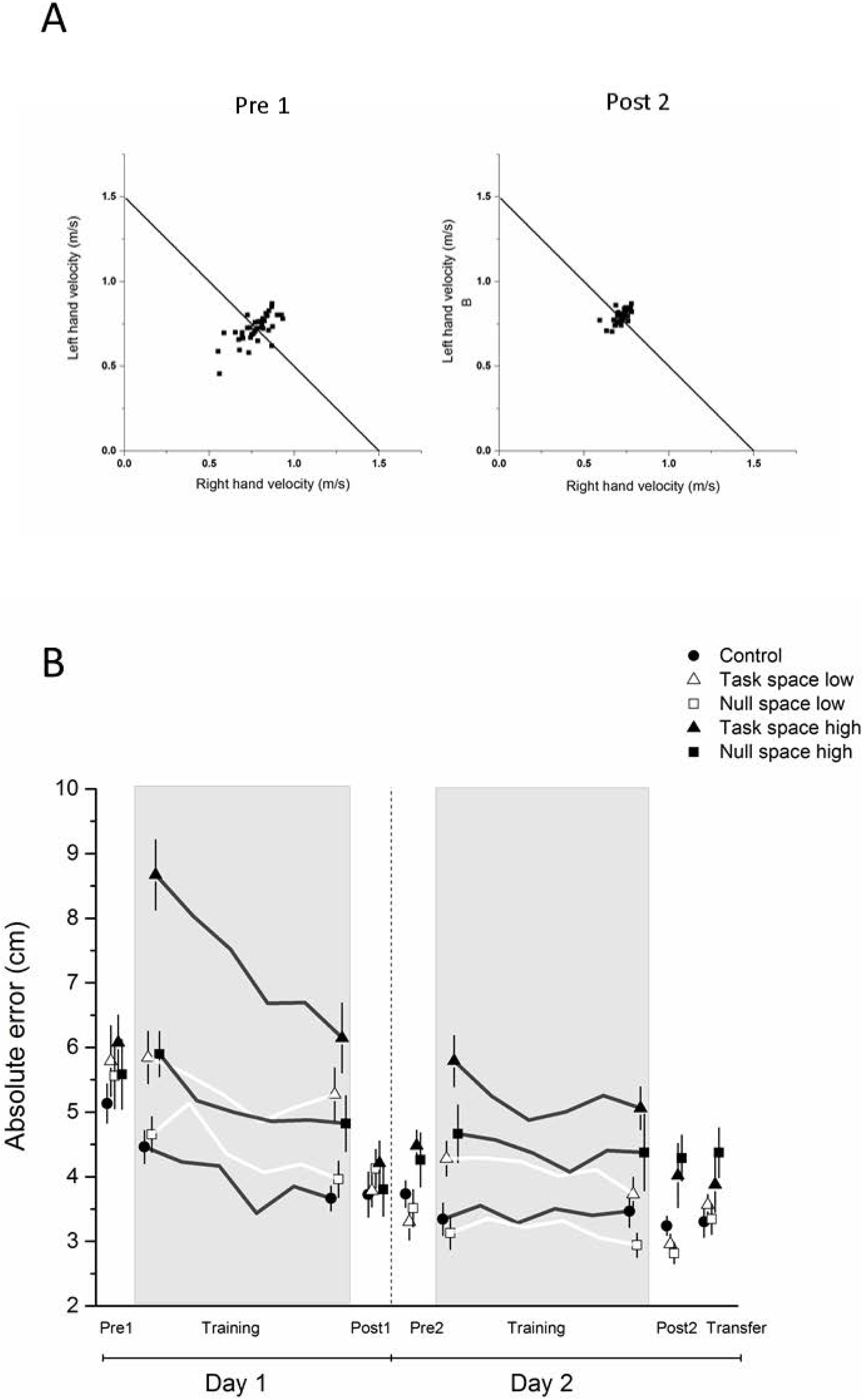
(A) Sample performance for one participant in the control group plotted in the velocity space at the start of training (pre-test 1) and at the end of training (post-test 2). Solid line indicates GEM, where there is zero task error. (B) Absolute error for all groups across both days of practice. All groups decreased absolute error with practice, but the high variability groups (task space high and null space high) had higher errors at the end of training. Training blocks (where different perturbations were applied to different groups) are highlighted in grey, whereas test blocks (where all groups performed in identical conditions) are shown with a white background.

Post hoc analysis of the interaction indicated that in pre-test1, there were no significant differences between the groups, but at post-test2, there was a significant effect of group – the null space high group had significantly higher absolute error than the control group. (p =.043)

For the contrast analysis of low variability and high variability groups, comparisons indicated no significant differences at pre-test1, but the low variability groups had significantly lower absolute error than high variability groups at post-test2, p =.001.

For the contrast analysis of the null space vs task space groups, comparisons were not significant either at pre-test 1 or post-test 2 (ps> 0.05).

### Task space variability

Task space variability, like absolute error, decreased with practice, but the improvements differed by group (Figure 4A). There was a significant main effect of test, F(3,126) = 62.18, p <.001, that was mediated by a significant test × group interaction, F(12,126) = 2.41, p =.007. The main effect of group was not significant, F(4,42) = 1.276, p =.295.

**Figure 4.**
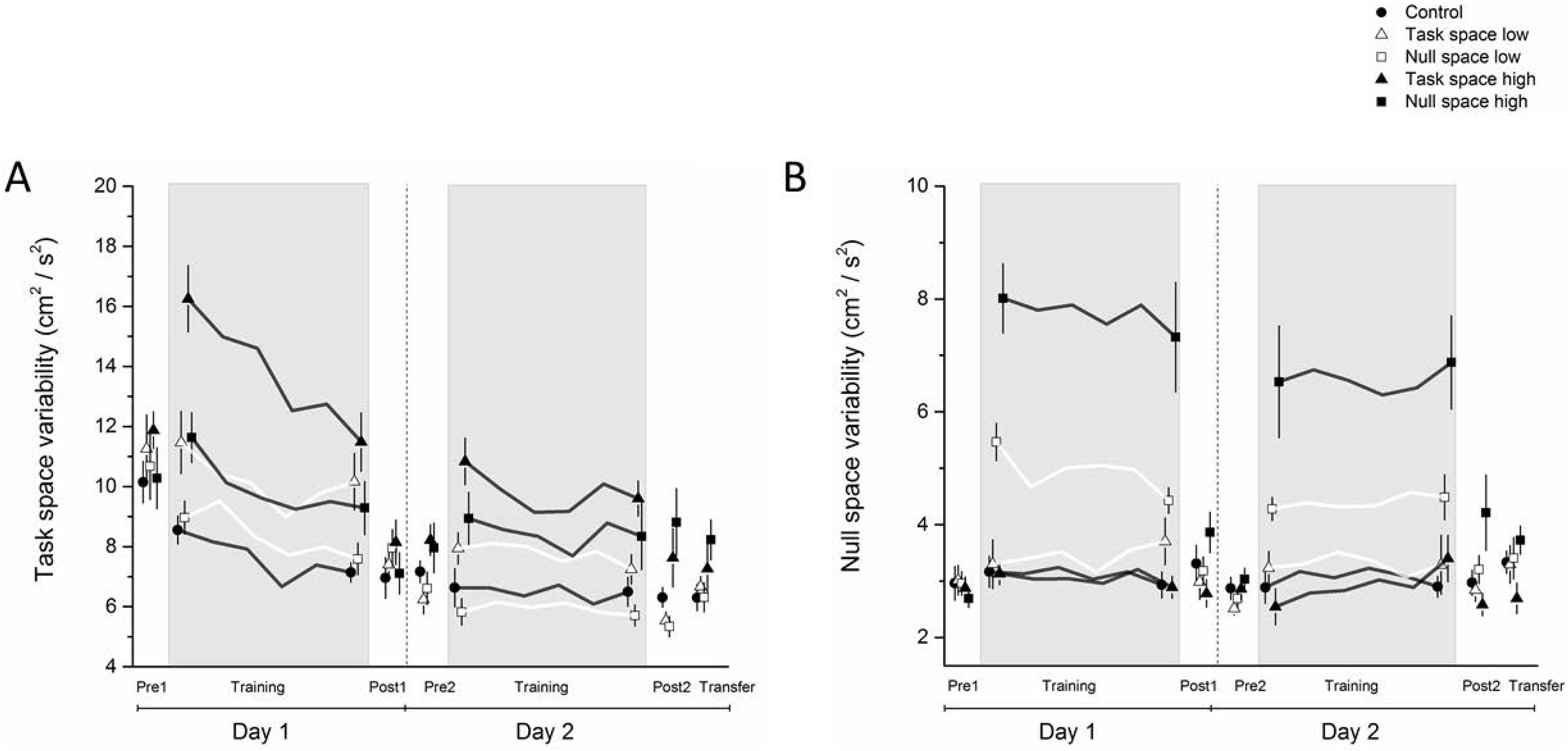
(A) Task space variability, and (B) Null space variability for all groups across both days of practice. Both task and null space variability decreased with practice for all groups. For task space variability, the high variability groups showed higher task space variability than the low variability groups in post-test2. For the null space variability, the null space groups showed higher null space variability thank the task space variability groups in post-test2. Training blocks (where different perturbations were applied to different groups) are highlighted in grey, whereas test blocks (where all groups performed in identical conditions) are shown with a white background.

Post hoc analysis of the interaction indicated that in pre-test1, there were no significant differences between the groups, but at post-test2, there was a significant effect of group – the null-space high group had significantly higher absolute error than the control group (p =.044).

For the contrast analysis of low variability and high variability groups, comparisons indicated no significant differences at pre-test1, but the low variability groups had significantly lower task space variability compared to the high variability groups at post-test2, p <.001.

For the contrast analysis of the null space vs task space groups, comparisons were not significant either at pre-test1 or post-test2.

### Null space variability

Null space variability, like absolute error and task space variability–decreased with practice, but the improvements differed by group (Figure 4B). There was a significant main effect of test, F(2.32,97.21) = 4.094, p =.015, that was mediated by a significant test x group interaction, F(9.26,97.21) = 2.13, p =.032. The main effect of group was not significant, F(4,42) = 1.881, p =.132.

Post hoc analysis of the interaction indicated that in pre-test1, there were no significant differences between the groups, but at post-test2, there was a significant effect of group – the null space high group had significantly higher null space variability than the control group (p =.050).

For the contrast analysis of low variability and high variability groups, comparisons indicated no significant differences at pre-test1, or post-test2.

For the contrast analysis of the null space vs task space groups, comparisons were not significant at pre-test1, but the null space groups had significantly higher null space variability than the task space groups in post-test2 (p =.007).

### Aspect ratio

The analysis of aspect ratio showed a greater reduction in the task space variability compared to the null space variability– all participants increased their aspect ratio with practice, but the improvements differed by group (Figure 5). There was a significant main effect of test, F(2.54,106.65) = 42.498, p <.001, and a main effect of group F(4,42) = 2.695, p =.044, that was mediated by a significant test × group interaction, F(10.16,106.65) = 2.64, p =.006.

**Figure 5.**
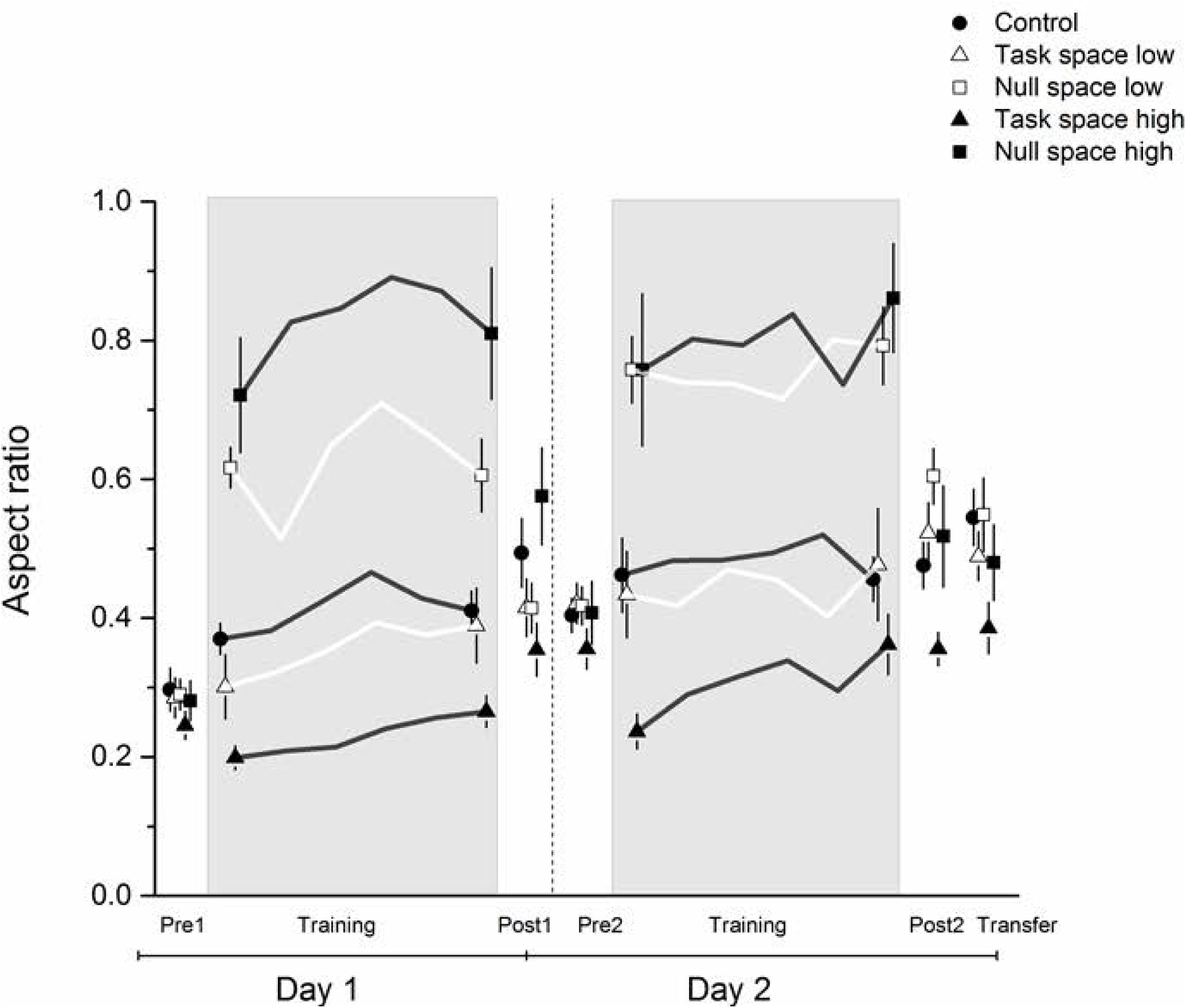
Aspect ratio (ratio of null to task space variance) for all groups across both days of practice. Introducing task and null space variability created differences in aspect ratios between groups during training, but these differences between null and task space groups also persisted at the end of training. Training blocks (where different perturbations were applied to different groups) are highlighted in grey, whereas test blocks (where all groups performed in identical conditions) are shown with a white background.

Post hoc analysis of the interaction indicated that both in pre-test1 and post-test2, there were no significant differences between any of the groups and the control group.

For the contrast of low variability and high variability groups, comparisons indicated no significant differences at pretest 1, but that the low variability groups had higher aspect ratio than the high variability groups (p =.007).

For the contrast analysis of the null space vs task space groups, comparisons were not significant at pre-test1, but the null space groups had significantly aspect ratio than the task space groups in post test 2 (p =.011).

### Autocorrelation in task and null space

For ACF-1 in task space (Figure 6), there was a significant main effect of test, F(3,126) = 6.948, p <.001, where the ACF-1 is post-test2 was greater (less negative) compared to pre-test1. The main effect of group and test × group interaction were not significant. None of the planned comparisons were significant.

For ACF-1 in null space, there were no significant main effects of test, group, or a test x group interaction. None of the planned comparisons were significant.

**Figure 6.**
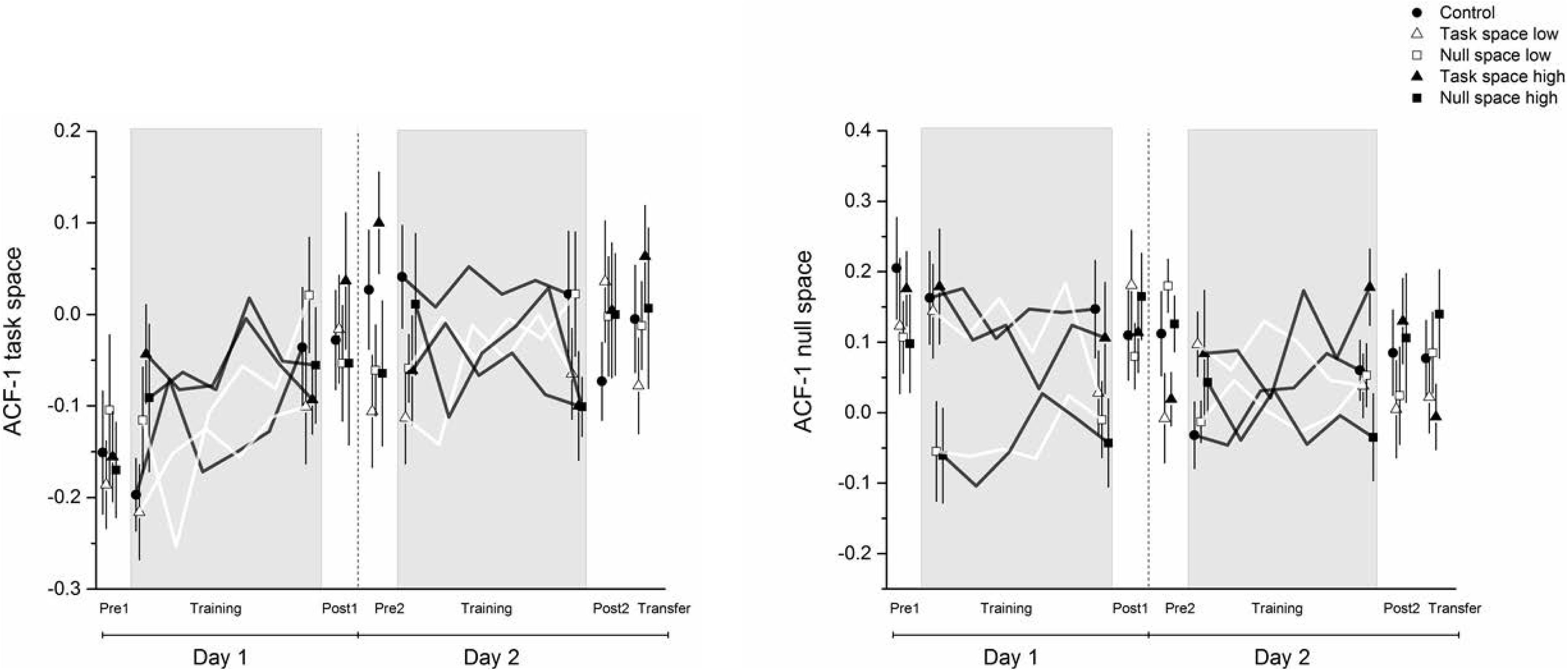
Lag-1 autocorrelation (ACF-1) in the task and null space for all groups across both days of practice. ACF-1 in the task space started negative early in learning and became closer to zero at the end of practice, whereas the ACF-1 in the null space started positive and did not change significantly with learning. Training blocks (where different perturbations were applied to different groups) are highlighted in grey, whereas test blocks (where all groups performed in identical conditions) are shown with a white background.

### Transfer test

In the transfer test, the results were similar to post-test 2. For the absolute error, there was a trend towards a main effect of group (F(4,42) = 2.168, p =.089). Post-hoc comparisons indicated that the null-high group had significantly higher error than the control group. Planned comparisons indicated the low variability groups had significantly lower error than the high variability groups (p =.032), but there was no statistical difference between the null space groups and the tasks space groups.

## Discussion

We examined the effect of introducing variability during the learning of a bimanual shuffleboard task. We introduced different amounts of variability in the null space and in the task space, and examined its effects on learning. We found that all groups were able to reduce the error with 2 days of practice. However, we found that the amount of variability introduced had a much larger influence on learning compared to the dimension in which the variability was introduced.

Specifically, introducing small amounts of variability had no (or even slightly positive) effects on learning compared to the control group where no variability was introduced, whereas introducing larger amounts of variability negatively affected learning regardless of whether the variability was introduced in the null or task space.

### General changes with learning – amount and structure of null and task space variability

We first describe the general changes in the null and task space variability with learning that were observed in all groups. As expected, all groups reduced absolute error and task space variability with learning. With respect to null space variability, there was a decrease with learning, but the reduction of variability in the null space was smaller than in the task space (measured as an increase in the aspect ratio). These results are consistent with the idea that while reduction in variability happens both in task and null space (i.e. people choose to be more consistent overall), variability in the null space reduced at a slower rate compared to the task space. Although the relative reduction of null and task space variability has been mixed in learning studies (Latash, 2010), these results support the hypothesis that learning in redundant tasks involves selective reduction in certain components of movement variability rather than a global reduction of overall variability (Abe and Sternad, 2013; Sternad et al., 2011; Yang and Scholz, 2005).

In addition to the amount of variability, we also examined the temporal structure (i.e. the trial to trial changes) in the variability using the lag-1 autocorrelation. The autocorrelations in the task space started off negative and went to zero with practice for all groups. In contrast, the autocorrelations in the null space were positive, and did not significantly change with learning. Overall, using a within-subject learning design, our results support and extend earlier studies (van Beers et al., 2013b; Dingwell et al., 2010), which showed that the task and null space not only differ in the amount of variance that is allowed to accumulate, but also in the trial-to-trial corrections.

### Effects of introducing null and task space variability on learning

With respect to group differences, the analysis of the absolute error clearly indicated that groups with high variability (whether it was null space and task space) showed greater errors compared to the groups with low variability. Practice with higher amounts of variability resulted in higher errors, that were associated with higher task space variability. In fact, the transfer test also showed a similar pattern of results (although the statistical significance was weaker), indicating no benefit to generalization relative to the control group.

Why would training with increased variability hurt motor learning? First, the fact that null and task space variability produce similar results on learning makes it unlikely that these are due to any mechanisms that are dependent on learning from error or exploration. In our view, the most likely explanation for these results is that learning in these tasks is use-dependent or model-free learning (Diedrichsen et al., 2010; Haith and Krakauer, 2013). Use-dependent learning occurs from repetition, where there is a tendency to make successive movements similar to each other (Jax and Rosenbaum, 2007; Ranganathan and Newell, 2010a). Introducing trial-to-trial variability (either in null or task space) may hamper the ability to make these similar movements from trial-to-trial and disrupt this learning mechanism to stabilize a solution.

Indeed, the discrepancy in effects of variability in prior studies can be linked directly to the learning task. Studies that have found beneficial effects of variability typically use adaptation paradigms (Singh et al., 2016; Wu et al., 2014), where participants need to find a new solution (e.g. moving the hand along a different trajectory) to be successful at the task. In these cases, variability is critical for exploration and finding a new solution. However, in the task used in the current study, participants did not have to necessarily find a novel solution, but rather reduce the variability around a desired solution. This learning mechanism is distinct from adaptation tasks and potentially also involves different neural substrates (Shmuelof et al., 2012). In these cases where reduction of variability is the critical factor to learning, increasing exploration in the task or null space may not be beneficial to learning. The current results are in agreement with prior studies that found that introducing variability in a precision aiming task did not facilitate learning relative to the control group (Ranganathan and Newell, 2010b, 2010c). Moreover, studies that have been successful in reducing variability employ strategies where task difficulty is increased without perturbing the actual movement itself – either through visual error augmentation, i.e. where errors appear bigger visually (Hasson et al., 2016) or by changing the reward structure (Huber et al., 2016).

### Dimension in which variability was introduced shaped movement strategy

Although the introduction of variability in the task and in the null space had similar effects on task performance, there were striking differences in the strategy employed by these groups. The null space groups showed much higher aspect ratios than the task space groups, indicating that practicing with more variability in the null space biased their strategy toward using more null space variability. These results support the notion that training with variability, even if not directly benefitting performance, still influenced the movement strategy that participants used in the task. These findings are consistent with recent studies showing how introducing variability during practice can be used to shape coordination in the learning of redundant motor tasks (Thorp et al., 2016), and could have direct applications in movement rehabilitation, where movement strategy is critical.

In summary, we found that motor variability influenced learning – but the amount of variability was more critical to learning than the dimension in which variability was introduced. These results support the idea that variability is a complex construct that has multiple ways of affecting motor learning (He et al., 2016; Ranganathan and Newell, 2013), and that careful consideration of the task demands is necessary to understand how variability may affect learning both in terms of the task outcome and movement coordination used to perform the task.

